# Object Detection Techniques for Live Monitoring of Amoeba in Phase-Contrast Microscopic Images

**DOI:** 10.64898/2026.03.30.715415

**Authors:** Olga Chambers, Ashley Cadby

**Affiliations:** The Department of Physics and Astronomy, The University of Sheffield, Hounsfield Rd, S3 7TH

**Keywords:** Machine learning, amoeba detection, deep-learning, object detection, phase-contrast image

## Abstract

In contemporary bio-imaging-based research, computer-based assessment is becoming crucial for the characterisation of biological structures, as it minimises the need for time-consuming human annotation, which is prone to human error. Furthermore, it allows for the use of optical techniques that use lower photon intensities, thereby reducing reliance on high-intensity excitation and mitigating adverse effects on their activities. This study details the development and evaluation of sophisticated deep-learning models for amoeba detection using phase-contrast imaging. Using a single-class annotated dataset comprising 88 images and 4,131 annotations, we developed nine object detection models based on Detectron 2 and six variants based on YOLO v10. The diversity of the dataset, acquired under varying setup parameters, facilitated a comprehensive evaluation of the strengths and limitations of each model. A comparative analysis of speed and accuracy was performed to identify the most efficient models for real-time detection, providing critical insights for future microscopic analyses.

## Introduction

The integration of artificial intelligence (AI) has become a cornerstone of modern biomedicine, revolutionising diagnostic precision in fields ranging from oncology to microbiology. By automating complex tasks, such as image segmentation and feature extraction, AI-driven methodologies significantly mitigate manual workloads and accelerate the analysis of pathological conditions. These advancements can be particularly transformative when applied to phase-contrast imaging for the detection of amoebas, which act as a model for human immune cells. By providing scalable and precise identification of cellular features, AI-enhanced imaging addresses long-standing diagnostic challenges, ultimately bolstering our capacity to monitor and respond across clinical and environmental contexts.

Despite the reported success of machine learning in various studies, predicting the most effective model for a specific task remains challenging. The 2018 Data Science Bowl, hosted by Kaggle, Booz Allen Hamilton, and the Broad Institute, challenged participants to advance state-of-the-art nucleus segmentation. The top-performing deep learning-based methods utilized a select few architectures, including Mask R-CNN, U-Net, and feature-pyramid networks (1). The U-Net (2, 3) is a widely used deep learning approach across numerous applications. Its architecture has various implementations tailored to the quality of datasets and research objectives. Although U-net can provide fast and accurate segmentation, its sensitivity to inhomogeneity and poor contrast between objects make it difficult to use in microscopic image processing without post-processing (3, 4). Moreover, dense cell populations pose challenges for many tasks. Cells may be in contact with each other, making it difficult to segment them from one another, and the level of difficulty increases as the population density increases. To overcome this limitation, researchers have used U-net in combination with object detection techniques, such as Fast R-CNN and YOLO(5). Thus Yi et al. (6) constructed a joint network based on the combined single shot multi-box detector (SSD)(7) and U-net techniques. The Faster R-CNN method has been used in low-contrast microscopic imaging (8). It has mainly been applied to analyse images with negligible cell overlaps and a small number of objects. Another well-known object detection model that has proven to work well is RetinaNet (9, 10). It uses a focal loss function to address class imbalance during training. Rahman at al. (11) indicates that it performs better than other well-known techniques for weed detection.

The motivation of this research is to compare some known object detection techniques in terms of accuracy and speed to improve amoeba segmentation and live tracking in phase-contrast images. This should help reduce fluorescence light usage for amoeba labelling during tracking and provide more accurate morphological feature extraction. Thus, Faster R-CNN, RetinaNet, and YOLO were selected based on a literature review and our previous experience.

### Amoebas are a model system for phagocytosis

Object detection techniques have several characteristics, including the number of model stages. Generally, techniques are categorised into one- and two-stage categories. The literature indicates that two-stage methods provide better accuracy; however, they have a higher computational cost (12). Two-stage detectors divide the object detection task into two stages: region-of-interest extraction, followed by feature extraction, bounding box regression, and classification. A single-stage detector ignores the region-of-interest extraction process and performs direct candidate processing.

Detectron2 (13) includes an open source high-quality implementations of state-of-the-art object detection algorithms such as Faster R-CNN and RetinaNet showing promising results in numeric applications (11, 14–17). It was implemented in PyTorch and was able to provide fast training on single or multiple GPU servers. Here, only models trained with a 3 × schedule (three times the number of epochs) were selected for comparison, as the 1 × models were heavily undertrained. Detectron2 provides object detection baselines based on three different backbone combinations, which were included in the comparative analysis. In contrast to YOLO and RetinaNet, Faster R-CNN is a two-stage object detection method. Seven Faster R-CNN architectures were tested with backbone (layer) depths of 50 and 101.

You Only Look Once (YOLO) is a popular object detection architecture known for its high speed and accuracy in many applications. It was first presented in a research paper by Redmon *et al*. at the Conference on Computer Vision and Pattern Recognition (CVPR) in 2016 (18) and was later implemented as an open-source algorithm. Although it has gained popularity in the computer vision community, some researchers have indicated that the YOLO method has difficulties distinguishing overlapping objects and positioning bounding boxes, which could be crucial for some medical problems, such as blood cell analysis (19). In this study, we used the latest extension, YOLOv10 (20). It is a one-stage object detection approach with multiple universal properties that improves upon previous versions. YOLO uses various model scales to cater to different application requirements. YOLOv10 has five variants, ranging from nanoscale to extra-large models. The choice of these models is based on the tradeoff between the required accuracy and inference time. Speed is of significant importance when determining the points scored on a shooting card.

Both Detectron2 and YOLO have been used in microscopy image processing; however, their popularity depends on the specific requirements of the task and the research community. In microscope data analysis, the choice between Detectron2 and YOLO depends on the specific requirements of the analysis. Detectron2 typically offers higher accuracy because of its use of two-stage detectors and flexibility to incorporate various backbone networks and feature pyramids. This is particularly important in microscope data analysis, where objects of interest may have complex shapes, low contrast, or require precise segmentation for accurate measurement and classification. Although YOLO is generally considered to be slightly less accurate than Detectron2, it still achieves impressive results, especially with newer versions, such as YOLOv4 and YOLOv5, which have incorporated improvements to enhance accuracy. For microscopy data, if the objects are well-defined and the task does not require pixel-level precision, YOLO can be an effective tool. The authors of (21) indicated that YOLO provides better and more meaningful detection of objects in a scene. However, Detectron2 provides a higher detection speed than YOLO, resulting in faster overall detection and tracking processes.

When working with relatively small datasets, incorporating pre-trained “backbones” (22) can be a strategic choice to ensure successful network training (23). The term “backbone” refers to the feature-extracting network, many of which have been pre-trained on the COCO dataset (24). Intuitively, more uniform data would result in more accurate model training. A comparative analysis between YOLO v10 and Detectron2 for object detection was performed using a dataset consisting of images acquired at different times with different parameters, resulting in large variability in the field of view, artefacts, and amoeba population. The comparison included pre-trained models that were included in the packages at the time of testing.

The remainder of this paper is organised as follows. Section 2 describes the research dataset and network architecture. In Section 3, we discuss the results of object detection using the selected techniques. We summarise our study and highlight the key conclusions in Section 4.

## Materials

### Datasets

The research dataset consisted of phase-contrast images of the amoeba *Dictyostelium discoideum* acquired using a Nikon Tie microscope equipped with a 60 × Plan Fluor Apo Ph3 objective lens (Nikon Instech, Tokyo, Japan). A Photometrics 95B sCMOS camera was used with an exposure time of 100 ms.

Data were collected at different times under various conditions. Some examples of the images are shown in 1. It can be seen that all images have similar structure where amoebas are near the same color as background. Owing to the various environmental conditions, the images have different artefacts. appearance. The bright “halo” surrounding cell borders and the “shade-off effect” artefacts produce low contrast inside the cells (25).

The image width varied between 1024 and 2048 pixels. The image with a circular field of view had the largest resolution (i.e., 2048 × 2808 pixels). One set of images contained yeast appearing as bright small circles forming stacks. It can be noted that several subsets of images contain a significant number of amoebas inside the field of view.

### Annotation

The acquired images were annotated for object detection and segmentation using only one class (i.e., “amoebas”). Polynomial annotation was performed manually using the online platform, “Make Sense” (26), which allows the export of annotations in the COCO JSON format that automatically generates box coordinates based on the polygon. Polynomial annotation helps minimise the impact of the background during training and validation. The resultant annotation file contains information about the source image file name, each amoeba polynomial coordinates in *x, y*, box coordinates in *x, y*, and associated class as region attributes. Detectron 2 can use this file for both, segmentation and object detection. For the YOLO model training, the data annotations were saved in the YOLO format for each image. **Training/validation set**. A research dataset was used to create two sets: training and validation. Table 1 provides information about the number of images and their corresponding annotations. For training and validation, we included images from all subsets collected during the research To investigate the impact of shading variations on object detection models, we adjusted the contrast of the images by saturating the bottom 20.

**Table 1.** Information about training and validation sets used for object-detection training.

| set | num. images | num. annotations |
| --- | --- | --- |
| training | 64 | 3156 |
| validation | 24 | 975 |

A personal computer (Dell, Inc., Round Rock, TX, USA) with a 13th Gen Intel (R) Core (TM) i7-13650HX CPU (2.60 GHz), 32.0 GB memory, and GeForce RTX 4060 GPU (Nvidia Co., Santa Clara, CA, USA) was used for training and final detection.

## Method

In the object detection procedure for the custom dataset described herein, four modules are involved: obtaining annotated datasets for training and validation, selecting the model configuration, and training and evaluation using appropriate metrics. All object detection models were implemented using open-source software packages and included available backbones for feature extraction that were pre-trained on the Common Objects in Context (COCO) dataset (24). The parameters that enabled the best performance for each technique were selected and are listed in Table 2. The remaining hyperparameters followed the default settings in the original implementation.

**Table 2.** Metrics definition.

| Metric | Abbreviation | Description |
| --- | --- | --- |
| Intersection Over Union | IoU | The overlap between ground truth and predicted bounding box |
| Average Precision | AP | Summary of the Precision-Recall Curve area under the curve for selected class/category |
| Average Precision at IOU=0.50 | AP@0.50 | Summary of the Average Precision at fixed IOU for selected class/category |

Neither framework has a strict maximum image size for training. However, there are practical limitations based on the hardware, specifically GPU memory. Larger images require more memory to store the input data, intermediate activations, gradients, and model parameters during training. The image sizes in the research dataset varied between 512 and 2048 pixels. Therefore, to balance the training process, a fixed maximum training size of 1024 pixels was selected. This means that when, an image with at least one side larger than 1024 pixels is rescaled to maintain the original proportion. The frameworks transform the predicted bounding box coordinates to fit the dimensions of the original image after the detection process. Additionally, the appropriate parameters corresponding to the overlapping rate between the detected objects in each method were changed to ensure that the detected mask corresponded to only one object, which is crucial for the tracking task.

The Ultralytics open-source implementation (**?**) was used for the YOLO evaluation. The Detectron2 library (13) was used to train the RetinaNet and Faster R-CNN models. The Detectron2 model configuration in this study had three components that were used to implement seven Faster R-CNN and two RetinaNet models with a 3 × learning rate scheduler.

- Backbone: ResNet (R) or ResNext (X)
- Backbones combinations: ResNet +Feature Pyramid Network (FPN), ResNet conv4 backbone with conv5 head (C4) or ResNet conv5 with dilations (DC5)
- Number of layers: 50 or 101

The FPN (27) is a well-known structure in object detection that fuses a shallow feature map and an upsampled deep feature map to detect objects of different scales. C4 was the original baseline in the Faster R-CNN framework.

Traditional model metrics were used to estimate the validation accuracy. Intersection over Union (IoU) is a common metric used to measure the amount of overlap between the ground truth (i.e., human annotation) and the predicted model boundary boxes. If the prediction is perfect, the IoU is equal to 1, and if it completely misses, the IoU is equal to 0. This means that the ground truth would be considered similar to that with an IoU of 0.50. The average precision (AP) provides deeper information about the model performance by using information on the positive and negative guesses of the model (28). Generally, a prediction with an IoU greater than 0.50 is considered a true positive prediction. The equations used to calculate these metrics are as follows:

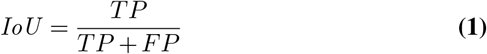

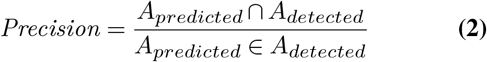

where *A* is the number of pixels within the box area, *TP* indicates that the predicted and manual annotations are the same, and *FP* indicates that the manual annotations were not predicted. Both *TP* and *FP* are calculated using a fixed IOU of 0.50.

The actual time difference in training these models depends on the dataset size, image resolution, hardware, and other factors. Thus, a comparison of was performed on a computer using cloth as a possible training parameter, such as batch size and number of workers. Table 3 lists the parameters that were fixed for training each model.

**Table 3.** Training hyperparameters.

| Hyperparameters | Detectron2 | YOLOv10 |
| --- | --- | --- |
| Learning rate | 0.00025 | 0.00025 |
| Max iterations | 1000 | 1000 |
| max image size | 1024 | 1024 |
| probability threshold | 0.5 | 0.5 |
| batch size | 4 | 4 |
| number of workers | 4 | 4 |

## Results and Discussion

In this section, we present the details of the comparative analysis of amoeba segmentation using Detectron2 and YOLOv10 in terms of accuracy and speed. We conducted the experiments on a personal computer (Dell, Inc., Round Rock, TX, USA) with a CPU (13th Gen Intel(R) Core(TM) i7-13650HX, 2.60 GHz), 32.0 GB of memory, and a GPU (GeForce RTX 4060; Nvidia Co., Santa Clara, CA, USA).

Tables 5 and 4 summarise the overall accuracy and speed of the object detection techniques for the validation set. Twenty-four random images were selected from the research dataset to evaluate the performance of the trained models and compare the output images with expert annotations. Detection was performed for two slices with sizes of 2048 × 2048 pixels containing 135 amoebas and 1200 × 1200 pixels containing 137 amoebas.

**Table 4.** Performance of the amoeba detection in phase-contrast space using YOLO library.

| Model architecture | early stop [iterations] | image@4128 [sec] | image@1200 [sec] | AP@0.50 |
| --- | --- | --- | --- | --- |
| YOLOv10 - 10n | 380 | 0.35 | 0.27 | 86.24 % |
| YOLOv10 - 10s | 316 | 0.47 | 0.39 | 87.63 % |
| YOLOv10 - 10m | 213 | 0.71 | 0.65 | **88.03 %** |
| YOLOv10 - 10b | 307 | 0.76 | 0.69 | 87.23 % |
| YOLOv10 - 10l | 260 | 0.94 | 0.88 | 87.45 % |
| YOLOv10 - 10x | 288 | 1.17 | 0.92 | 86.02 % |

**Table 5.** Performance of the amoeba detection in phase-contrast space using Detectron2 library.

| Model architecture | early stop [iterations] | image@4128 [sec] | image@1200 [sec] | AP@0.50 |
| --- | --- | --- | --- | --- |
| Faster RCNN +R50-C4 | 1000 | 0.69 | 0.59 | 88.1% |
| Faster RCNN +R50-DC5 | 1000 | 0.53 | 0.42 | **89.41 %** |
| Faster RCNN +R50-FPN | 1000 | 0.5 | 0.41 | 88.75% |
| Faster RCNN +R101-C4 | 1000 | 0.71 | 0.56 | 88.22% |
| Faster RCNN +R101-DC5 | 1000 | 0.52 | 0.4 | 88.78% |
| Faster RCNN +R101-FPN | 1000 | 0.5 | 0.38 | 88.62% |
| Faster RCNN +X101-FPN | 1000 | 0.47 | 0.42 | 88.52% |
| RetinaNet + R50 | 1000 | 0.49 | 0.47 | **86.78%** |
| RetinaNet+R101 | 1000 | 0.5 | 0.49 | 86.78% |

The tables present the training times for various object detection models at different image sizes. The models include several variants of YOLOv10 (yolov10s, yolov10m, yolov10n,yolov10l, yolov10b, and yolov10x), Faster RCNN with different backbones (R50-C4, R50-DC5, R50-FPN, R101-C4, R101-DC5, R101-FPN, and R101-XPN), and RetinaNet with different backbones (R50 and R101).

Here is an observation of the results shown in the table:

- YOLOv10 reached early-stop criteria before processing maximum defined iterations when Detectron2 used all defined iterations for model training in each case.
- The results obtained during comparative analysis does not indicate significant difference between the performance of Detectron2 and YOLO models accuracy. The mAP50 scores of these models range from 84 %to 89% that can be explained not just miss-segmentation but over-segmentation when other elements such as yeast were identified falsely as amoeba.
- Among the YOLOv10 variants, the yolov10n model has the fastest training time for both image sizes, while the yolov10x model has the slowest training time. For the Faster RCNN models, those with the FPN (Feature Pyramid Network) backbone generally have faster training times compared to those with C4 or DC5 backbones, especially for the smaller image size.
- Larger image sizes result in more pixels and, consequently, more computational work for the model. This also lead to longer training times per epoch since the model has to process more data. Larger images typically provide higher-resolution features, which may be beneficial for detecting small objects. This can potentially lead to better accuracy, particularly if the task involves identifying fine details or small objects within the images.
- R101 is deeper than R50, with 101 layers compared to 50 layers. This indicates that R101 has a higher capacity to learn complex features, which can lead to better detection accuracy, particularly for challenging datasets with varied and complex scenes. R50 is less complex and requires fewer computations than the R101 model. This results in faster feature extraction and, consequently, faster training and inference times. For real-time or near-real-time applications where speed is critical, R50 might be preferred.
- The RetinaNet models have relatively fast training times, with the RetinaNet+R50 being slightly faster than RetinaNet+R101 for the larger image size, but the difference is negligible for the smaller image size.
- All Detectron2 models outperformed the YOLO models. The best-performing Detectron2 model is the Faster RCNN + R50-DC5, with an AP@50 score of 89.41%. YOLO v10 models also performs good, how-ever with less acuuracy. YOLOv10m achieved the highest AP@50 value of 86.03%.

If inference time is the critical requirement in a real-time environment, YOLOv10s and Faster R-CNN +R50-FPN in Detectron2 would be better choices.

To provide a visual inspection of the models on the test dataset, six images from the test dataset are randomly chosen and are shown in Figure For each library, the models that showed the best results in terms of accuracy were selected. Notably, all three models exhibited good results, accurately identifying and localising objects within the images.

Figures 2 shows the results of amoeba detection using different models.

**Fig. 1.**
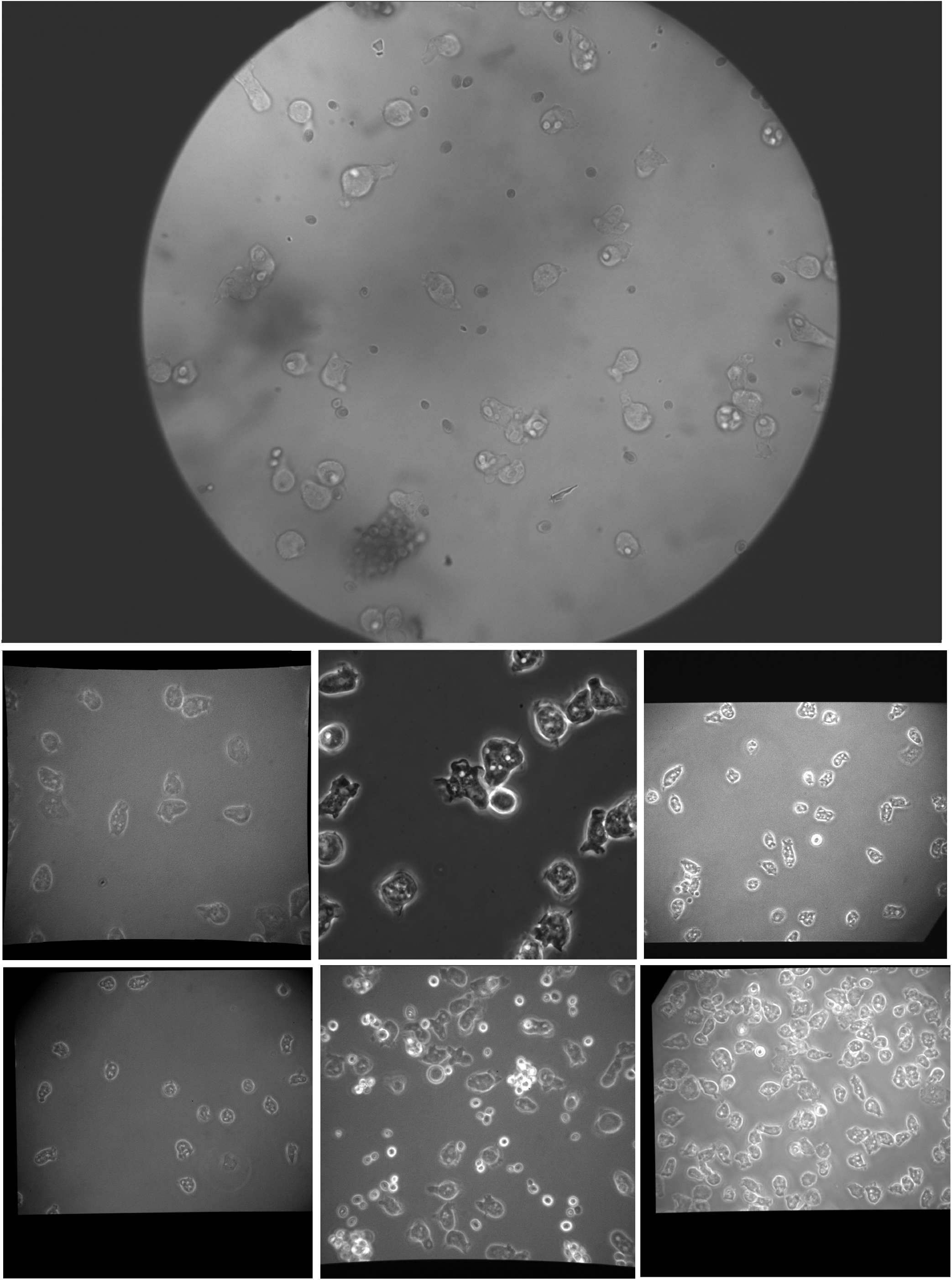
Examples of images used in this work

**Fig. 2.**
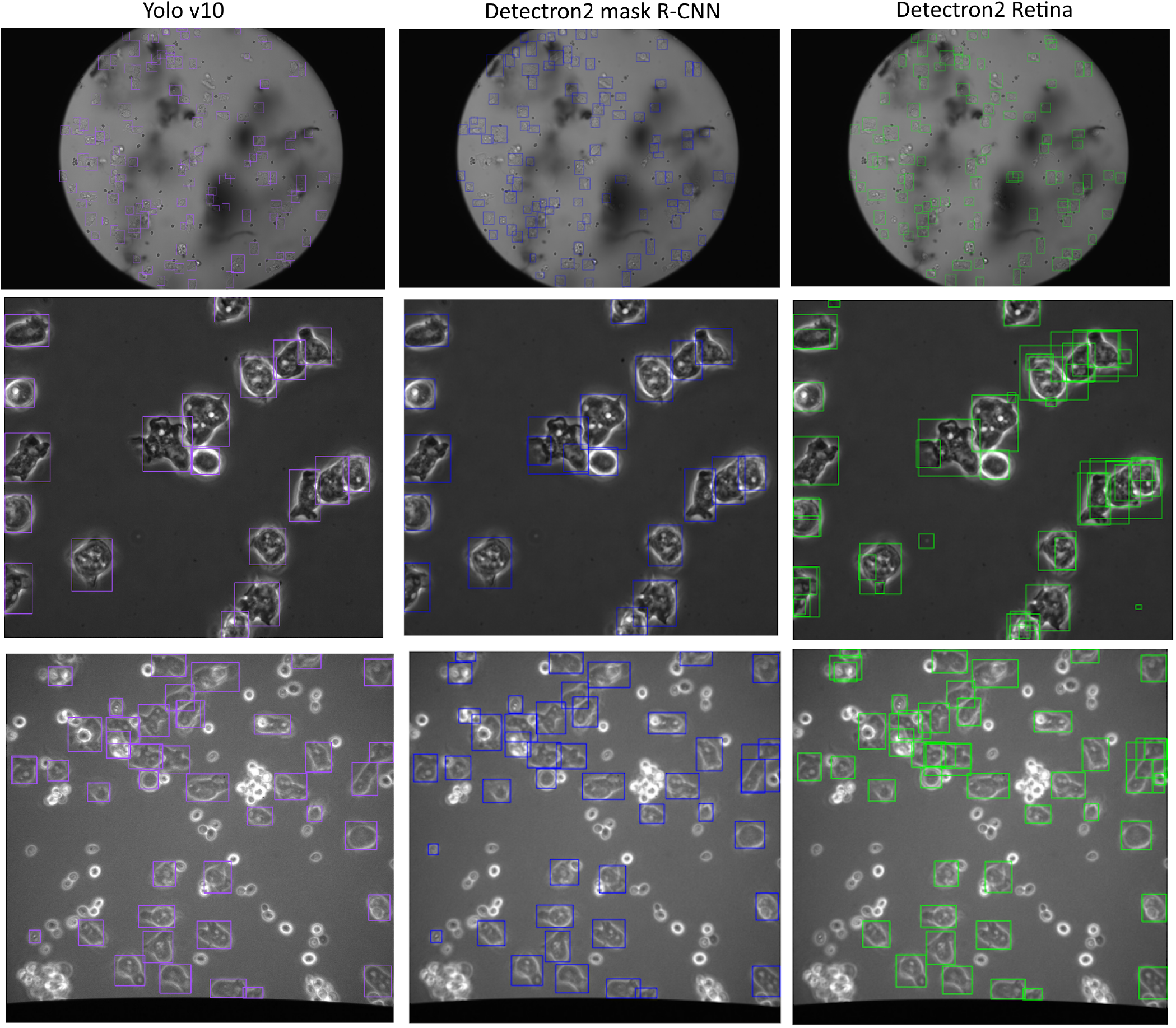
Amoeba detection result in randomly selected phase-contrast image using different techniques

Concerning the depth of the network, it is expected that the changing the depth from 50 to 101 increases the quality of the results. Models with deeper or larger backbone networks yield higher detection accuracies. However, our results show that a larger backbone does not always cause better quality but always diminishes the speed owing to higher computational complexity.

Among the three Faster R-CNN configurations, those employing DC5 achieved the best accuracy performance with a relatively high speed. Therefore, we considered this backbone for amoeba segmentation in phase-contrast images. Faster R-CNN (R101-C4), despite its longer inference time, did not achieved the highest overall detection accuracy.

Although each of Detectron2 and YOLO has strengths and weaknesses, overall we observed that Detectron2 returned more meaningful results than YOLO. In many cases, Detectron2 detected each amoeba separately, whereas YOLO detected them multiple times, thereby reducing accuracy and making object tracking more challenging.

Both methods exhibited good performance in images with inhomogeneity at the field-of-view edges.

Detectron2 Faster RCNN +R50-FPN model displays a faster tracking detection than YOLOv8 and produces more accurate detections when there is a limited dataset for training. On the contrary, the original configuration of the Faster R-CNN employing a conv4 (C4) backbone is shown to be the slowest among the assembly that also conform with reported in similar study (29).

YOLO resulted in a lower accuracy rate than Faster R-CNN, which performed over-segmentation more often than other techniques. Nevertheless, YOLO showed the potential for real-time deployment because of its fast inference, which achieved a comparable detection accuracy of 85.63 Figure shows an improved result of YOLOv8 when training is performed for the same research dataset but excluding images with circular field of view (see Figure). Nevertheless, we present the performance of the techniques with a dataset that combines different appearances that are more common in real life and provide more accurate characterisation of the model performance stability. Data augmentation through geometric transformations can improve the accuracy of weed detection. During amoeba detection, we faced the problem of detecting multiple overlapping boxes corresponding to the same amoebae. Unfortunately, these boxes have similar probabilities; therefore, they cannot be simply thresholded. Additional post-processing would clarify this; however, the criteria that select boxes with higher accuracy significantly increase the computational time of the overall process, even when the original detection is fast. In this context, Faster R-CNN and EfficientDet were the most efficient, whereas RetinaNet generated the most overlapping boxes.

Another notable difference between YOLO and Detectron usage is that YOLO does not require any model parameters to be defined, apart from the pre-trained weights, whereas Detectron2 requires configuration parameter definition to match the pre-trained weights.

In general, YOLO is often chosen for its speed in terms of both training and inference. However, Detectron2, being a more flexible and feature-rich framework, might offer advantages in terms of accuracy and customisation, which could justify longer training times for certain applications. This finding was confirmed in the present study.

Note that the latest versions of both YOLO and Detectron2 continue to evolve, and their performance may vary with each new release.

In this study, we compared several neural network architectures designed for object detection. We believe that this comparison will be valuable to researchers developing amoeba detection strategies and promoting the effectiveness of machine learning in biomedical analysis.

There is often a trade-off between the speed of a model (in terms of inference time) and its accuracy. Larger images can yield more accurate models; however, they come at the cost of slower inference times. Smaller images can speed up the process; however, they might lead to lower accuracy, especially if important details are lost due to downscaling. Training with smaller images might help the model generalise better if the dataset is limited, as it forces the model to learn more abstract features rather than details that might not be relevant.

## Conclusion

In this study, we report the results of our comparative analysis of common object detection methods for amoeba allocation in phase-contrast images under different conditions. The pretrained Faster R-CNN, RetinaNet, and YOLOv10 were finetuned using a research dataset that comprised images with a non-uniform structure. The obtained results help to understand the strengths and weaknesses of the models and their sensitivity to image structural variations. Study shows that Faster R-CNN slightly over-performs Yolo. Nevertheless, all methods showed relatively similar results and had potential for use in biomedical tasks. The results were obtained using a relatively small dataset; therefore, the reported results could potentially be improved. We believe that the obtained results can help to understand the potential of these methods for real-time tasks in biomedical applications.

